# Single Particle Tracking of Genetically Encoded Nanoparticles: Optimizing Expression for Cytoplasmic Diffusion Studies

**DOI:** 10.1101/2024.11.17.623896

**Authors:** Elizaveta Korunova, Vitali Sikirzhystki, Jeffery L Twiss, Paula Vasquez, Michael Shtutman

## Abstract

Single particle tracking (SPT) is a powerful technique for probing the diverse physical properties of the cytoplasm. Genetically encoded nanoparticles provide an especially convenient tool for such investigations, as they can be expressed and tracked in cells via fluorescence. Among these, 40-nm GEMs provide a unique opportunity to explore the cytoplasm. Their size corresponds to that of ribosomes and big protein complexes, allowing us to investigate the effects of the cytoplasm on the diffusivity of these objects while excluding the influence of chemical interactions during stressful events and pathological conditions. However, it has been shown that cytoplasmic viscosity is tightly regulated and plays a crucial role in maintaining homeostasis during protein synthesis and degradation. Despite this, the effects of GEM expression levels on diffusivity remain largely uncharacterized in mammalian cells. To optimize the GEMs tracking and estimate GEMs-expression effects we constructed dox-inducible GEM expression system and compare with a previously reported constitutive expression system. The optimized level of GEMs expression increases the measured diffusivity from 0.29 ± 0.02 μm^2^/sec in GEMs-overexpressed cells to 0.35 ± 0.02 μm^2^/sec; improve homogeneity throughout the cell population; and facilitates particle tracking. We also improved the analyses of GEM diffusivity by applying effective diffusion coefficient while considering the type of motion and assessing the heterogeneity in the type of motion by calculating the standard deviations of particle displacements.

**Statement of significance:** Describing cytoplasmic properties, such as environmental viscosity and protein complex motion, is essential for understanding molecular-level changes in cell function and pathology. A recently developed approach uses self-assembling fluorescent protein probes, expressed in cells, to investigate cytoplasmic properties through single-particle tracking (SPT). One such system employs genetically encoded multimeric (GEM) nanoparticles— scaffold protein structures similar in size to ribosomes. This study addresses a key limitation in SPT of GEMs by examining how varying GEM expression levels affect measured diffusivity and tracking quality in mammalian cells. Our findings demonstrate that controlled GEM expression reduces particle overcrowding, increases measured diffusivity, and enhances track detection. This work contributes valuable insights into optimizing GEM nanoparticle applications for studying cytoplasmic viscosity and motion dynamics.

## Introduction

The intricate spatial, compositional and dynamical complexity of the cytoplasm contributes to its varied physical properties [1]. Depending on size and speed—whether driven by molecular motors or thermal motion— constituents in the cytoplasm may experience it as a viscous fluid, viscoelastic material, poroelastic substance, or even glass-like material [2-6]. More specifically, larger organelles, such as mitotic spindles, nuclei, mitochondria, and large lysosomes, experience the cytoplasm as a viscoelastic or poroelastic medium because they perceive the cytoskeleton as a continuous structure. When diffusing within the cytoplasm, smaller objects like ions, metabolites, proteins, and protein complexes (e.g., lipoprotein complexes, proteasome, spliceosome, ribosomes) can experience varying effects based on their size. If significantly smaller than the major crowding elements (e.g., the cytoskeletal mesh, particularly actin, and ribosomes), these objects perceive the cytoplasm as a viscous medium. However, if their size is comparable to these major crowding elements, they may experience the cytoplasm as a viscoelastic medium. Studies have shown that the microrheological properties of the cytoplasm, such as elasticity and viscosity, are critical in regulating various cellular activities, including enzymatic [7] and metabolic activity [3], microtubule dynamics [8], cell division, differentiation, apoptosis [9], protein synthesis and degradation [10] and liquid-liquid phase separation transitions [11]. Subsequently, changes in microrheological properties accompany pathological processes. For example, an increase in cytoplasmic viscosity, correlated with elevated levels of the signaling molecule H2S, has been proposed as a biomarker for Parkinson’s disease [12]. Further, a fluorescent probe, designed with a metabolic molecule size and capable of penetrating the blood-brain barrier, has been proposed to target Golgi and detect increased viscosity in the early stages of Alzheimer’s disease [13]. Significant changes occur at the mesoscale, corresponding to the size range of protein complexes and organelles (10–1000 nanometers in diameter). For instance, a decrease in viscosity has been observed to accompany cellular enlargement during aging in fibroblasts [14]. An increase in diffusivity at mesoscale level has also been shown to be a necessary response to starvation, triggering the efficient formation of Q-bodies—membranous organelles that regulate the degradation of misfolded proteins during stress [11].

Microrheological properties of cytoplasm can be analyzed using various techniques. Bulk methods [15] measure viscosity as an average parameter among a cell population by tracking the rotational diffusion of probes, molecules designed to estimate these properties. Single-cell methods, such as fluorescence recovery after photobleaching [16], fluorescence decay after photoactivation [17], fluorescence lifetime imaging microscopy [18], fluorescence correlation spectroscopy [19], and single particle tracking [20], allow for viscosity estimation within individual cells. Among these, single particle tracking (SPT) is a useful approach for estimating the diffusive properties of biomolecules in cytoplasm. SPT is based on the analysis of tracks from moving particles. One common method for analyzing these trajectories is to calculate the mean squared displacement (MSD) of particles as a function of lag time (τ) [21-24]. For many biological fluids, the MSD exhibits a power-law relationship with lag time, MSD ∼ τ^α^. This lag-time dependence reveals three possible types of particle motion: sub-diffusive motion (0 < α < 1), which can be caused by factors such as obstacles, crowding, or correlated motion; Brownian motion (α = 1), representative of a random walk of unhindered particles; and super-diffusive motion (α > 1), often indicative of directed or persistent movement (Figure 1A) [23].

**Figure 1.**
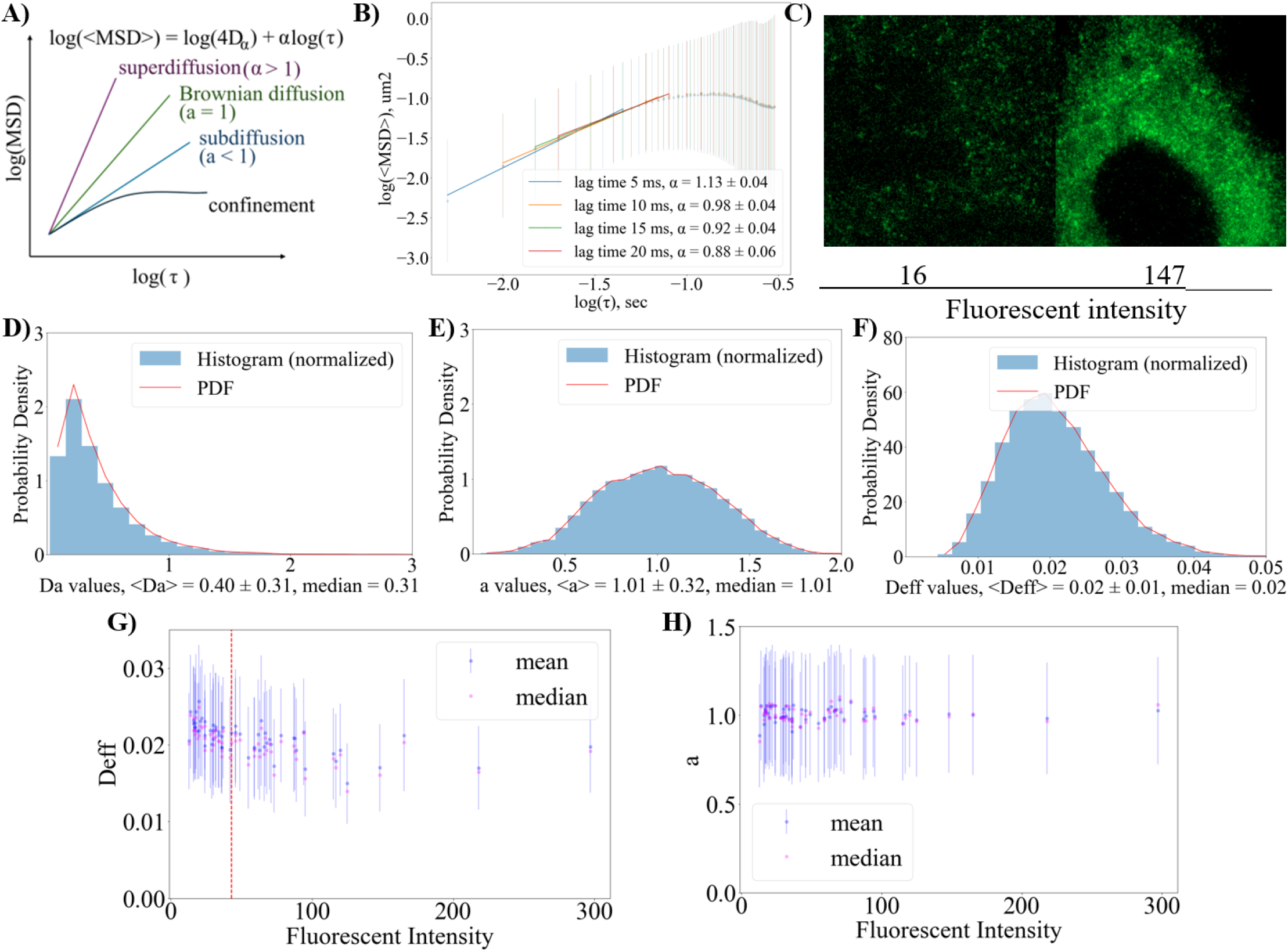
A) Characteristic log-log plots of MSD vs. lag time (τ). Slope α indicates diffusion type: Brownian (α=1), subdiffusive (α<1), superdiffusive (α>1). Plateau indicates confinement. B) Log-log plot of ensemble- and time-averaged MSD (<MSD>_ate_) for particles longer than 10 frames (50 ms), corrected for static error and fitted to a power law with different lag time Δt values of 5, 10, 15, or 20 ms (τ = nΔt), the corresponding alpha values are given in the legend; C) image of U2OS with lower (fluorescent intensity = 16 r.u.) and higher GEM expression (fluorescent intensity = 147 r.u.). Sample size was 60 cells. D) Density probability function of collected D_α_, E) α,and F) D_eff_ collected after fit with power-law relationship with σ/Δt = 0.5; G) Plot of D_eff_ vs. fluorescence intensity in cells throughout the U2OS colony. The red line indicates the division between cells with “High GEM expression” (30 cells) and “Low GEM expression” (30 cells); H) Corresponding plot of α vs. fluorescence intensity.

Previously, the particles had to be inserted into the cells by variety of physical techniques, and the difficulties in delivery restricted the application of the methods. The recent introduction of self-assembling nanoparticles with defined properties has revolutionized the field [25, 26]. Genetically Encoded Multimeric (GEM) nanoparticles for SPT are created by expressing scaffold proteins from the bacterium *Pyrococcus furiosus* decorated with the fluorescent protein Sapphire [25]. These scaffold proteins self-assemble into 40 nm nanoparticles, comparable in size to ribosomes and other big protein complexes, and may exhibit similar diffusive behavior in the cytoplasm, excluding chemical interactions. A significant advantage of this system is its ability to be expressed in eukaryotic cells, eliminating the need for microinjection and following cytoplasmic dilution. To date, GEM systems have been used to describe cytoplasmic viscosity in budding and fusion yeast, fungus, bacteria and mammalian cultures, such as HEK, A338 cell line, C. elegans and D. melanogaster, to study the effects of molecular crowding and spatial heterogeneity of cytoplasm [27-33] and even viscosity of nucleoplasm in HEK [34]. However, current analyses of GEMs do not account for varying levels of GEM expression in mammalian cells. These variations can impact the quality of SPT data, leading to issues such as particle overcrowding, inability to distinguish particles during SPT, and shorter detected tracks. Additionally, overexpression and, as a result, overcrowding of GEMs may influence the homeostasis of cytoplasmic protein concentration. For example, a recent study with *Xenopus egg* extracts demonstrated that cytoplasmic protein concentration is maintained through a negative feedback system based on viscosity. In this system, an increase in macromolecule concentration leads to increased cytoplasmic viscosity, which, in turn, stimulates higher rates of protein degradation relative to translation. This compensatory response pushes the system back toward its normal viscosity and macromolecule concentration [10]. Here, we investigate whether diffusivity values calculated from SPT analyses differ across varying levels of GEM expression and identify the optimal expression level for microrheological studies in mammalian cells. To optimize the GEMs tracking and estimate their expression effects, we constructed dox-inducible GEM expression system and compare with a previously reposted constitutive expression system. Our results show that GEM overexpression decreases measured diffusivity, while controlled GEM expression improves tracking. We also improved the analyses by applying the calculation of the effective diffusion coefficient while considering the types of motion and assessing the heterogeneity in motion types by calculating the standard deviation of particle displacements.

## Results

### The increase of measured GEM diffusivity accompanies the decrease in GEM expression

Assuming a power-law relationship, we analyzed GEM diffusivity in U2OS cells and optimized the analysis pipeline to correct the SPT data for localization errors introduced during imaging because improper correction can lead to false subdiffusivity (static error) or superdiffusivity (dynamic error; see the Methods section for details) [29, 35]. Briefly, we corrected ensemble- and time-averaged MSD (<MSD>_ate_), that was already corrected for static error, for dynamic error in two ways: following the criterion σ/Δt << 1 (Figure 1B) [35], where σ is acquisition time, and by directly removing the error (Figure S1) [29]. We then applied this analysis to single particle tracks, selecting only those consistent with power-law relationship (Figure 1A). This step is crucial to filter out particles exhibiting confined or stationary behavior that do not actively participate in motion. Such confined and stationary particles are commonly observed in cells. To compare particles of different α, we calculated each particle’s D_eff_ value (the ratio of its diffusion coefficient to diffusion coefficient in water) from the distributions of D_α_ and α (Figure 1D, F, E). We observed slightly higher diffusivity, with broader distributions for the D_eff_ coefficient, and α of approximately 0.95 when dynamic error was directly removed (Figure S1). In comparison, analyses using the criterion σ/Δt≪1 yielded an α closer to 1. Both approaches yielded results consistent with reported values for GEM diffusivity in U2OS cells. Based on these findings, the second approach, the criterion σ/Δt≪1, was selected for subsequent analyses. Summarizing, motion type was α ∼ 1 (Brownian), with ensemble-averaged diffusivity D_α_ ∼ 0.29 ± 0.02 μm^2^/s (σ/Δt = 0.5, table 1) and a median D_eff_ of 0.02 ± 0.01, corresponding to 0.31 ± 0.02 μm^2^/s after error correction.

**Table 1.** Microrheological parameters extracted from Figure 1 B and C.

| lag time | $D_a$, [$\mu\text{m}^2/\text{sec}^\alpha$] | $\alpha$ | $D_{\text{eff}}$ |
| --- | --- | --- | --- |
| 5.0 ms | $0.37 \pm 0.03$ | $1.13 \pm 0.04$ | $0.02 \pm 0.002$ |
| 10.0 ms | $0.29 \pm 0.02$ | $0.98 \pm 0.04$ | $0.019 \pm 0.001$ |
| 15.0 ms | $0.27 \pm 0.02$ | $0.92 \pm 0.04$ | $0.019 \pm 0.002$ |
| 20.0 ms | $0.25 \pm 0.02$ | $0.88 \pm 0.06$ | $0.019 \pm 0.002$ |
| dynamic error correction | $0.37 \pm 0.03$ | $0.97 \pm 0.02$ | $0.024 \pm 0.002$ |

To analyze GEM diffusivity in U2OS cells with varying GEM expression levels, we measured the mean fluorescence of GEMs in individual cells and calculated the distribution of D_eff_ for these cells. We then applied Spearman and Pearson correlation analyses to examine the relationship between the median D_eff_ and the mean GEM fluorescence in the cells (Figure 1G). We found a strong negative correlation for the measured median D_eff_, indicating a monotonic decrease (Spearman) in D_eff_ with increasing GEM fluorescence (Table 2). This was complemented by a moderate negative correlation (Pearson), suggesting an average linear decrease in D_eff_ as GEM fluorescence levels rose. We found no correlation between measured median α and GEM fluorescence levels, suggesting that while GEM expression levels influence diffusivity (Figure 1H), they do not affect their motion type in U2OS cells.

**Table 2.** Spearman (monotonic change) and Pearson (linear change) correlation coefficients for the dependency of median D_eff_ vs. fluorescence intensity.

| Experiment | Statistic | Value | p-value |
| --- | --- | --- | --- |
| power-law fit | Spearman Correlation | -0.71 | 3.04E-10 |
|  | Pearson Correlation | -0.56 | 3.47E-06 |

To further analyze the heterogeneity of α within cells, we examined the standard deviations of the x and y particle displacements. This approach was chosen because the classical MSD calculation introduces stochastic noise into the microrheological parameters (Figure S2), making it difficult to distinguish individual particle motion type accurately. We modeled the tracks of particles following standard Brownian motion (α = 1) and mixed (subdiffusive and standard Brownian) behavior (0.4 < α < 1) using a fractional Brownian motion model. Our analyses revealed that plotting the standard deviations of the x and y coordinates enables us to distinguish the heterogeneity of the system caused by change in motion type (Figure 2A and B). We found that U2OS cells population displays a certain degree of heterogeneity in particle movement between cells (Figure 2C). Notably, we observed minimal heterogeneity in α within cells (Figure 2D). Most cells exhibited behavior consistent with a uniform α throughout the cell.

**Figure 2.**
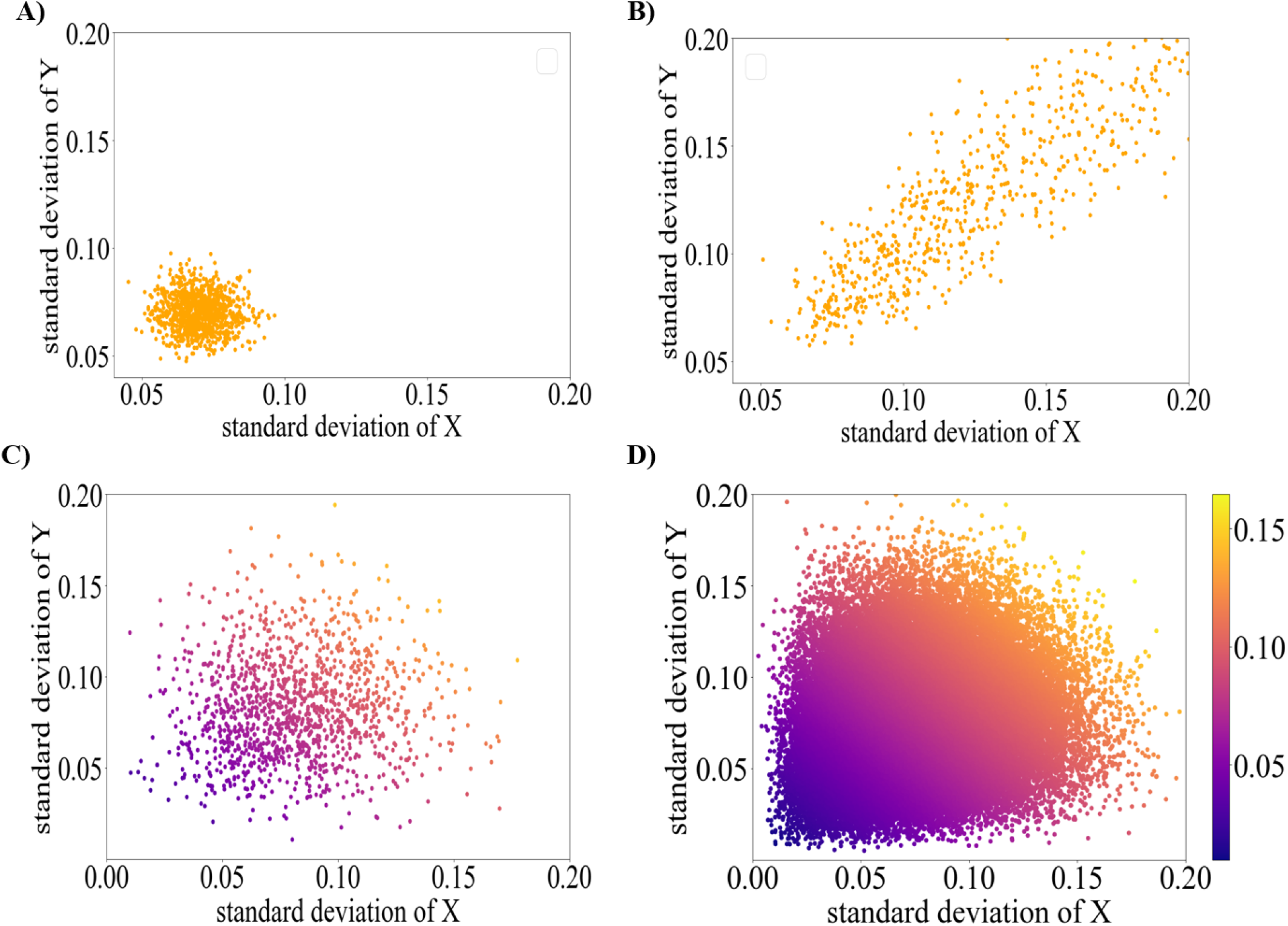
Plots of the standard deviation of x and y displacements at 10 ms intervals from the modeled data: A) Standard Brownian motion; B) Mixed behavior, combining subdiffusive and standard Brownian motion (0 < α < 1); Plots of the standard deviation of x and y displacements at 10 ms from experiment C) Plot for a single cell from the experiment; D) Plot of the standard deviation of x and y displacements throughout the colony.

## Controllable GEM expression in U2OS under doxycycline inducible promoter

To create a system with controlled GEM expression in mammalian cells, we developed U2OS cells expressing GEM under doxycycline induction at concentrations of 0.24, 0.5, and 1 μg/mL for 24 and 48 hours (Supplementary Videos S1, Videos S2, Videos S3, Figure S3). We processed the data as previously described and estimated the distribution of D_eff_ across cells based on their fluorescence levels (Figure S3). The Spearman analysis revealed no correlation between the measured diffusion parameters and fluorescence levels (Supplementary Table 1). To compare the measured diffusivity in the U2OS system with the uncontrolled expression of GEM, we categorized the system with the uncontrolled expression described above into “High GEM Expression” and “Low GEM Expression” groups (Figure 1 G, H, and I, Figure 2D, Supplementary Video S4 and S5). The threshold for division was determined by the fluorescence intensity achieved with doxycycline expression by some cells on the second day. To compare diffusivity among cells in the ‘High GEM Expression’ group, the ‘Low GEM Expression’ group, and U2OS colonies expressing GEM under doxycycline, we conducted a Bootstrap analysis. This analysis revealed that the ‘High GEM Expression’ group exhibiting lower diffusivity compared to the ‘Low GEM Expression’ group and compared to all colonies expressing GEM under doxycycline on the first day and certain colonies on the second day of expression (Figure 3A).

**Figure 3.**
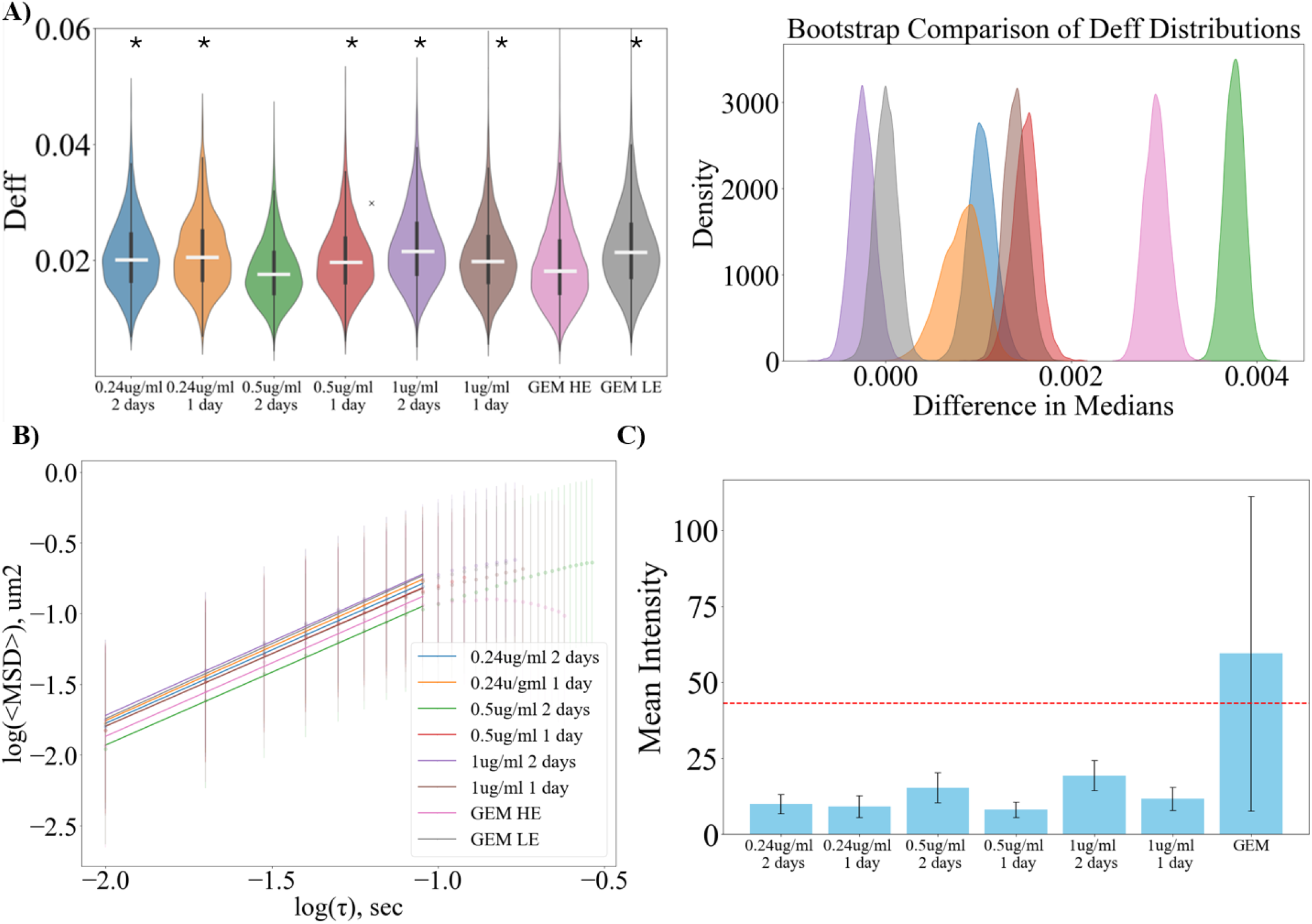
A) (left) Violinplots of D_eff_ distributions from experiments with different levels of GEM fluorescence (30 cells per group, except 0.24 ug/ml dox 1 day (20 cells) because of low expression) collected from power-law fit with σ/Δt = 0.5. GEM HE = ‘High GEM Expression’. GEM LE = ‘low GEM Expression’. ‘*’ significant difference with GEM HE estimated by non-parametric bootstrap analysis; (right) The corresponding bootstrap distributions reflecting median differences between groups; B) <MSD>_ate_ for particles passed single particle analysis for colonies expressing GEM under doxycycline induction, as well as for colonies with low and high expression of GEM without expression control. Vertical lines ∼ standard deviation; C) differences in mean fluorescence level (r.u.) between groups. The red line is the expression level between ‘High GEM Expression’ and ‘low GEM Expression’ groups.

We next analyzed ensemble and time-averaged diffusivity in the groups though calculating the <MSD>_ate_ of tracks from SPT analysis (Figure 3B). We found that the lower GEM expression increases measured diffusion coefficient D_α_ from 0.29 ± 0.02 um^2^/sec up to 0.35 ± 0.02 um^2^/sec (α = 1), which corresponds to an increase in D_eff_ from 0.019±0.002 to 0.021±0.001 (Table 2). Analyzying the heterogeneity of α within groups (Figure 4) showed clear heterogeneity in motion type for groups with higher GEM expression compared to those with lower GEM expression.

**Figure 4.**
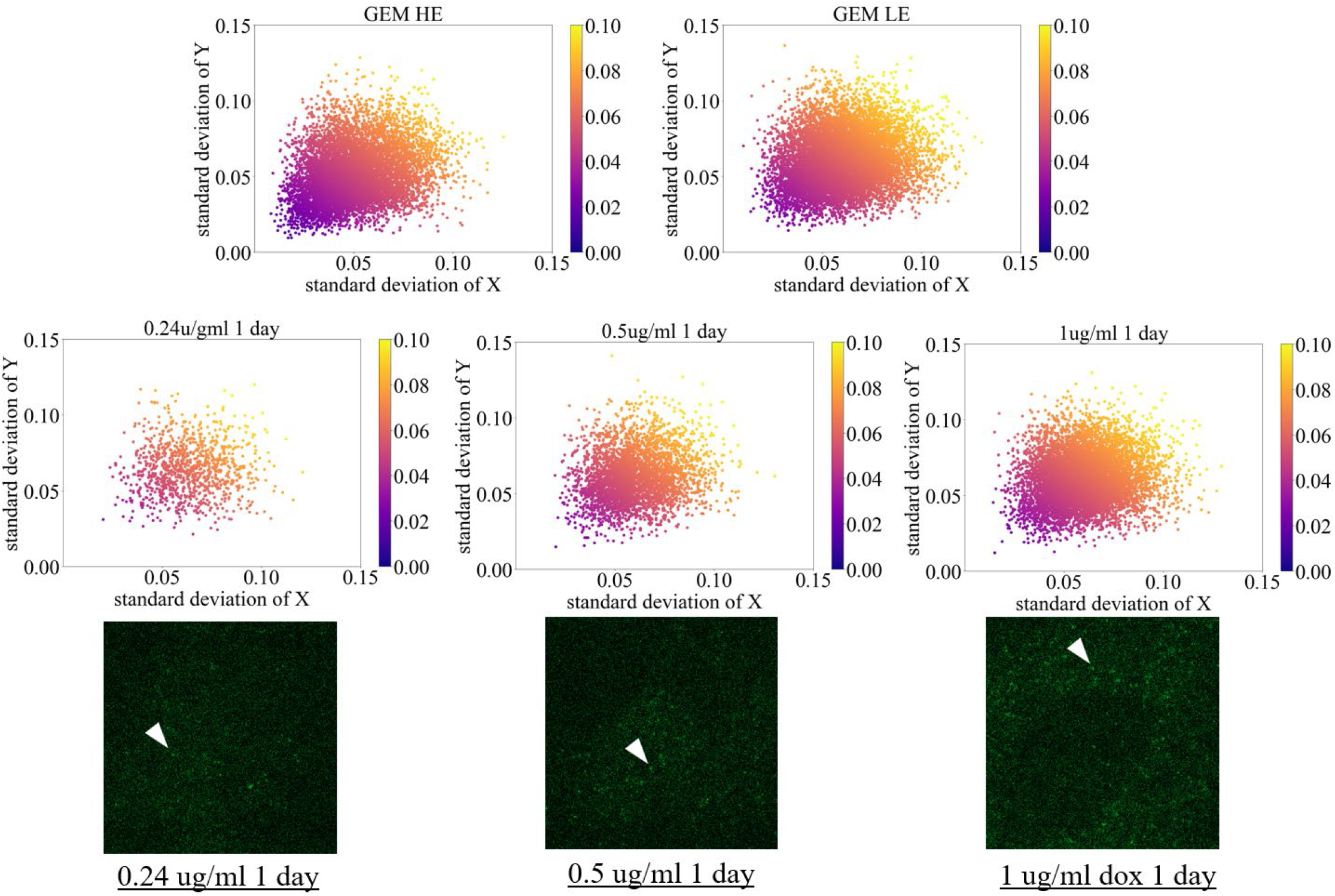
Plots of the standard deviation of x and y displacements at 10 ms for colonies expressing GEM under doxycycline induction for the first day and corresponding video (Video S1, S2 and S3) from single particle tracking, along with those exhibiting low and high expression of GEM without expression control.

## Discussion

In this study, we aimed to estimate the influence of GEM expression levels on diffusivity in mammalian cells. First, we evaluated the correlation of GEM diffusivity across a GEM-expressing cells population under a constitutive promoter. To do this, we applied a power-law fit to the MSD, corrected for static error and reduced dynamic error using established guidelines (σ/τ << 1 [35]). We found that averaged type of motion for particles thought U2OS colony corresponds to Brownian motion (α = 1), that corresponds with diffusivity described for 40 nm size in U2OS in previous reports [6, 11]. The estimated diffusion coefficient also corresponds to previously reported values for GEM in mammalian cells (Figure 1B, table 1) [11, 25, 30]. We then applied the Spearman correlation, which revealed that the Spearman coefficient indicates a strong monotonic decrease in the median measured diffusion coefficient (D_eff_) within the cell as the mean fluorescence of GEM (GEM expression) increases (Figure 1G, table 2). Notably, the difference in GEM fluorescence is clearly visible in live imaging (Figure 1C). Finally, we detected heterogeneity in α among U2OS cells expressing GEM under a constitutive promoter (Figure 2).

Next, we created a lentiviral vector to express GEM under doxycycline induction and measured diffusivity at varying doxycycline concentrations and incubation times. As a result, we received U2OS cells with lower expression of GEM (Figure 3C). We then compared the diffusivity of GEMs in the constitutive promoter group, divided into ‘High GEM expression’ and ‘Low GEM expression’ groups (Figure 1C, 1I) with the diffusivity if inducible GEMs (Figure 3A). Using non-parametric bootstrap statistics, we found a significant increase in diffusivity (based on confidence intervals, Figure 3A) in both the ‘Low GEM expression’ group and the doxycycline-induced groups incubated for 24 hours, compared to the ‘High GEM expression’ group (from 0.29 ± 0.02 um^2^/sec up to 0.35 ± 0.02 um^2^/sec (α = 1), table 3). Additionally, through heterogeneity analysis based on standard deviation, we found that an increase in GEM expression is associated with the emergence and increase of heterogeneity in α (Figure 4).

**Table 3.** D_a_, α, and D_eff_ extracted from <MSD>_ate_ (Figure 6).

| | $D_a \text{ um}^2/\text{sec}^\alpha$ | $\alpha$ | $D_{eff}$ |
| --- | --- | --- | --- |
| 0.24 ug/ml 2 days | $0.33 \pm 0.01$ | $1.02 \pm 0.02$ | $0.0204 \pm 0.0007$ |
| 0.24 ug/ml 1 day | $0.35 \pm 0.01$ | $1.04 \pm 0.01$ | $0.0209 \pm 0.0005$ |
| 0.5 ug/ml 2 days | $0.28 \pm 0.01$ | $1.02 \pm 0.02$ | $0.0174 \pm 0.0007$ |
| 0.5 ug/ml 1 day | $0.32 \pm 0.01$ | $1.01 \pm 0.02$ | $0.0198 \pm 0.0008$ |
| 1 ug/ml 2 days | $0.35 \pm 0.01$ | $1.03 \pm 0.03$ | $0.0217 \pm 0.0012$ |
| 1 ug/ml 1 day | $0.31 \pm 0.01$ | $1 \pm 0.03$ | $0.0198 \pm 0.001$ |
| GEM High Expression | $0.29 \pm 0.02$ | $0.99 \pm 0.06$ | $0.0185 \pm 0.0019$ |
| GEM Low Expression | $0.35 \pm 0.02$ | $1.04 \pm 0.04$ | $0.0213 \pm 0.0014$ |

These findings indicate that the level of expression influences the diffusivity of GEM itself, likely due to the intracellular crowding effects of the GEMs (Figure 1C). Furthermore, a decrease in GEM expression appears to enhance the detection of longer tracks (Figure S4) and improves particle tracking by reducing the likelihood of false detections caused by particles moving closely together. However, the dependency between the level of GEM expression of its diffusivity can impact other aspects of cellular biophysics. Reports suggest that the viscosity of cells is tightly regulated, with fluctuations in cytoplasmic viscosity playing a key role in controlling protein synthesis and degradation [10, 27, 36]. Thus, although the decrease in GEM expression was accompanied by an increase in diffusivity, compensatory mechanisms for changes in microrheological properties in mammalian cells may exist and are yet to be determined.

## Methods

### Plasmid construction

The plasmid containing pCMV-pfv-paGFP for the expression of photoswitchable GEM was constructed from pCMV-pfv-Sapphire-Ires-DsRed (Addgene #116934) and paGFP_UtrCH (Addgene #26738) using the NEBuilder® HiFi DNA Assembly Master Mix kit for recombinant cloning. pCMV-pfv-Sapphire-Ires-DsRed was digested with XmaI and MluI to obtain the vector. paGFP was amplified from paGFP_UtrCH with the following primers: forward (atctatttccggtgaattcctcgagatggtgagcaagggcgagg) and reverse (ttgattgttccagacgcgttcacttgtacagctcgtccatgcc) using the NEBNext Ultra II Q5 Master Mix to generate the insert for recombinant cloning.

The plasmid containing TRE-pfv-Sapphire for the doxycycline-inducible expression of GEM was constructed from pCMV-pfv-Sapphire-Ires-DsRed (Addgene #116934) and pLIX403_Capture1_APOBEC_HA_P2A_mRuby (Addgene #183901) using the NEBuilder® HiFi DNA Assembly Master Mix kit for recombinant cloning. pLIX403_Capture1_APOBEC_HA_P2A_mRuby was digested with NheI and BshTI to obtain the vector. Pfv-Sapphire was amplified from pCMV-pfv-Sapphire-Ires-DsRed using the following primers: forward (gatcgcctggagaattggctagcatgctctcaataaatccaaccc) and reverse (gtggtggtggaccggttcatttgtacaattcatcaataccatg) with the NEBNext Ultra II Q5 Master Mix to generate the insert for recombinant cloning.

### Virus production and U2OS transduction

To produce lentivirus with the vector of interest, 7 × 106 293FT cells were plated in 10 mL of media (DMEM with 10% FBS, 1% Penicillin-Streptomycin-Glutamine) on PEI-coated 15 cm dishes. The next day, 1 mL of transfection solution containing 150 mM NaCl, 0.5 μg/μL psPAX2, 0.2 μg/μL pDM2.G, and 6 μg of one of the vectors of interest (Addgene #116934, pCMV-pfv-paGFP, or TRE-pfv-Sapphire) was prepared and incubated for 20 minutes at room temperature before transfection. 6-8 hours after plating, the 293FT cells were transfected using this solution. At 24 hours after transfection, the medium was changed. Supernatants were collected at 48 hours (stored at 4°C) and 72 hours after transfection, filtered with a 0.45 μm PES filter, and then centrifuged at 53,000g at 4°C using a JA-25.5 rotor in a Beckman Coulter Avanti J-20XP for 18 hours. The virus was resuspended in 200 μL of PBS and stored in 20 μL aliquots at -80°C until use.

50-60% confluent U2OS cells were plated on 35 mm dishes and transfected the next day with the virus of interest (Addgene #116934, pCMV-pfv-paGFP, or TRE-pfv-Sapphire). U2OS cells expressing pCMV-pfv-Sapphire-Ires-DsRed (Addgene #116934) or pCMV-pfv-paGFP were sorted by flow cytometry using a Cell Sorter SH800S, while U2OS cells expressing TRE-pfv-Sapphire were selected with 2 μg/mL puromycin.

U2OS human osteosarcoma cells (American Type Culture Collection) were cultured in high-glucose Dulbecco’s Modified Eagle Medium (ATCC, Manassas, VA), supplemented with 10% fetal bovine serum (Mediatech Inc., Manassas, VA) and 1% penicillin/streptomycin (HyClone, GE Healthcare Life Sciences, Pittsburgh, PA). Cells were maintained at 37°C in a humidified atmosphere with 5% CO2.

### Live imaging of U2OS expressing GEM to extract SPT data

U2OS cells expressing pCMV-pfv-Sapphire-Ires-DsRed (Addgene #116934) or TRE-pfv-Sapphire were plated on Delta T Dish 0.17 mm Black (04200417B) at a confluency of 0.1×106 cells and imaged using a Zeiss LSM 700 microscope with 493/517 nm excitation/emission (U2OS cells with pCMV-pfv-paGFP were additionally activated with 405 nm). Imaging was performed at maximum speed, approximately 5 ms between frames (200 frames per second), at 35°C using a heating stage (Delta T5 Culture Dish Controller) and a 100x oil objective. U2OS cells expressing TRE-pfv-Sapphire were imaged after induction with doxycycline at 35°C for 25, 29, or 48 hours with concentrations of 1 μg/ml, 0.5 μg/ml, and 0.1 μg/ml. To validate the absence of diffusion during imaging with stage and estimate the static error, U2OS cells expressing pCMV-pfv-Sapphire-Ires-DsRed (Addgene #116934) plated on Delta T Dish 0.17 mm Black (04200417B) were fixed with 4% PFA and imaged as other samples.

### Microrheological analysis of SPT data

Images obtained after single particle tracking (SPT) were analyzed in Fiji (ImageJ) using MosaicSuite 2D particle tracking software (parameters: radius = 6, cutoff = 0, per/abs = 1, linking = 1, displacement radius = 5). Subsequently, tracks with a length equal to or greater than 10 frames (50 ms) were further analyzed using a custom Python scripts[37] to extract the diffusion coefficient of particles and the α parameter, which characterizes the type of particle motion [22].

Briefly, data from particle tracking (x, y, time) were interpolated (5 ms, approximate acquisition time) to facilitate calculations and absence of drift was validated from velocity distribution for 5 ms. The time-averaged mean squared displacement (<MSD>_te_) and ensemble- and time-averaged mean squared displacement (<MSD>_ate_) were computed for different lag times, τ = nΔt, where n=1,2,3… length of track. To fit <MSD>_te_ or <MSD>_ate_ we designed a python code to estimate the linearity of log(<MSD>) vs log(τ) graph with treshold of fitting based on R^2^. The graph log(<MSD>_te_ **or** <MSD>_ate_) vs log(τ) was fitted to a power-law form, corresponding to anomalous diffusion [23]:

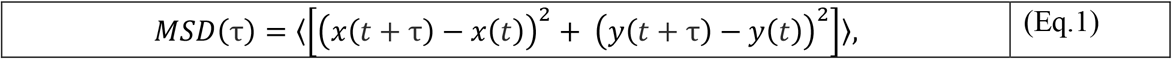

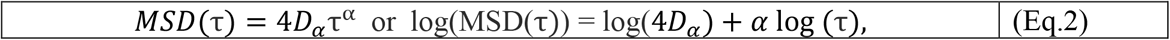

As the power law parameter α approaches 1, the movement resembles Brownian motion, where particles experience continuous, random fluctuations. Values below 1 lead to subdiffusive motion, where particles take smaller jumps and spread slower than Brownian motion. Conversely, α exceeding 1 signifies superdiffusive motion, with particles taking larger jumps and exploring space much faster.

Static localization error, mainly from inaccuracies in estimating the Point Spread Function (PSF) center during particle tracking— introducing artificial subdiffusive behavior—was corrected by subtracting the static MSD (calculated from a fixed sample) from the MSD, following the method of Savin and Doyle [35].

To estimate direct dynamic localization error we implemented an alternative correction approach, previously applied to GEM-expressing systems in E. coli (Eq.3) [29]:

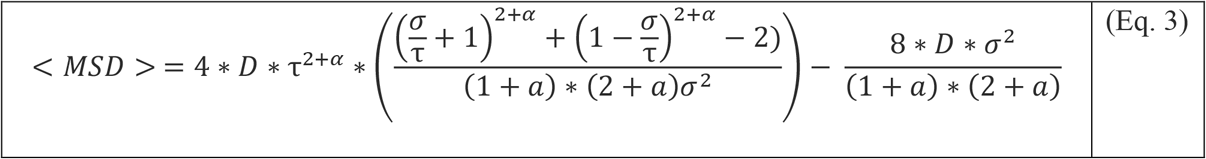

Alternatively to the direct correction of dynamic error, we reduced the influence of dynamic error by adhering to the criterion σ/Δt << 1, where σ represents an acquisition time, and Δt is the time interval for lag time (figure 1 B: τ = nΔt, where lag time Δt = 5, 10, 15, or 20 ms) [35].

We evaluated both direct error correction and the adherence to the σ/Δt << 1 criterion approaches to <MSD>_ate_. The fitting process was automatically halted if the fit accuracy dropped below an R^2^ value of 0.99.

We then applied an analysis with localization error correction to the pool of <MSD>_te_ values collected from individual tracks. We subtracted static <MSD> from <MSD>_te_ and tested the fit to a power-law function with ⧍t = n10ms, (σ/Δt = 0.5) and with direct dynamic error correction as described above, adjusting only the R^2^ filter threshold to 0.9 because of lower accuracy of <MSD>_te_. We also filtered out tracks where parameters of diffusion were more than five times higher than their corresponding errors. However, we found that the number of particles exhibiting power-law behavior without direct diffusion estimation, and with high accuracy, is greater than those following the power-law form after direct correction (Figure S2). Thus, we decided to use the power-law fit with σ/Δt = 0.5 for further estimation of diffusivity.

Finally, since the diffusion coefficient, D_α_, has units that depend on the power-law exponent α, this makes it difficult to compare the behavior of different particles and different regions within the cell. To overcome this challenge, we normalize the diffusion coefficient by the diffusion coefficient of GEMs in water,

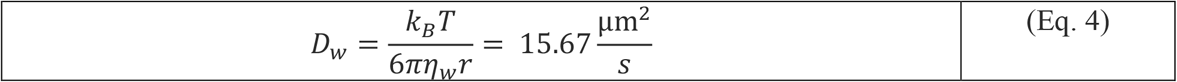

where the thermal energy is *k*_*B*_*T =* 4.11 pN nm, the viscosity of water is *η*_*w*_= 0.7195 mPa·s for T = 35 °C, respectively[38], and GEMs radius is *r =* 20 nm.

The effective diffusion coefficient is non-dimensional and given by [39],

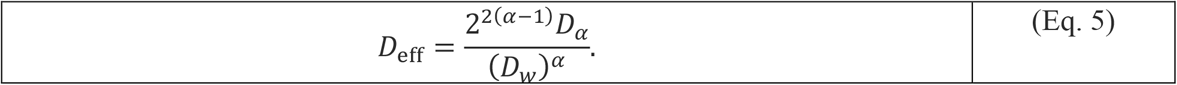

### Heterogeneity Analysis

To perform a heterogeneity analysis, we modeled particle movement using fractional Brownian motion, implemented it through Python scripting[37] and based the analysis on the standard deviations of particle displacements.

## Author contributions

EK, VS, JLT, PV and MS initiated the study, EK, VS, MS designed and performed experiments, EK, VS, MS and JLT analyzed the data and provided critical suggestions, EK, VS performed image analysis, EK and PV developed analytical tools, EK, VS, JLT, PV and MS wrote the manuscript.

## Acknowledgements

We thank COBRE Center for Targeted Therapeutics, Microscopy and Flow Cytometry Core for image analysis and Functional Genomics Core for transcriptomics analysis. The COBRE CTT cores are supported by NIH NIGMS P20GM109091. The work was supported by awards from NIH NIDA R21DA058586, R01DA054992, NIH NIGMS P20AGM109091-10S3 (to MS), NSF-DMS-1751339 (to PV), R01NS117821, R01NS089633, and the Dr. Miriam and Sheldon G. Adelson Medical Research Foundation (to JLT). JLT is the incumbent SC SmartState Chair in Childhood Neurotherapeutics.

