## Supplemental Figures for "Single Particle Tracking of Genetically Encoded Nanoparticles: Optimizing Expression for Cytoplasmic Diffusion Studies"

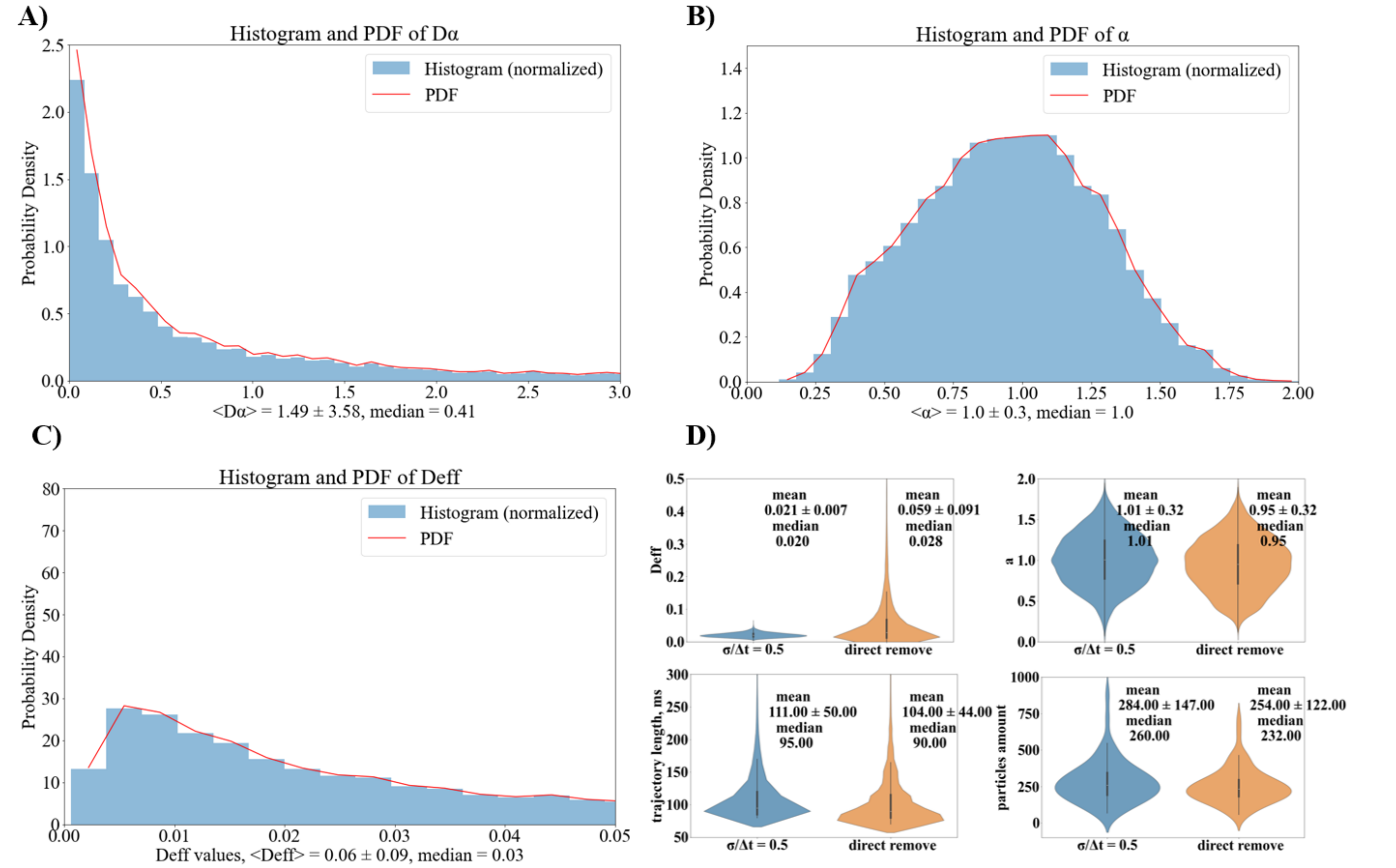


S1. A) Density probability function of collected $D_\alpha$; B) Probability distribution of α values collected after fitting with the power-law function, including dynamic error correction; C) Density probability function of collected $D_{eff}$ from power-law fitting with dynamic error correction. D) Violin plots of $D_{eff}$, α, trajectory length, and particle count from analyses using the power-law function (σ/Δt=0.5 ) and direct dynamic error correction.

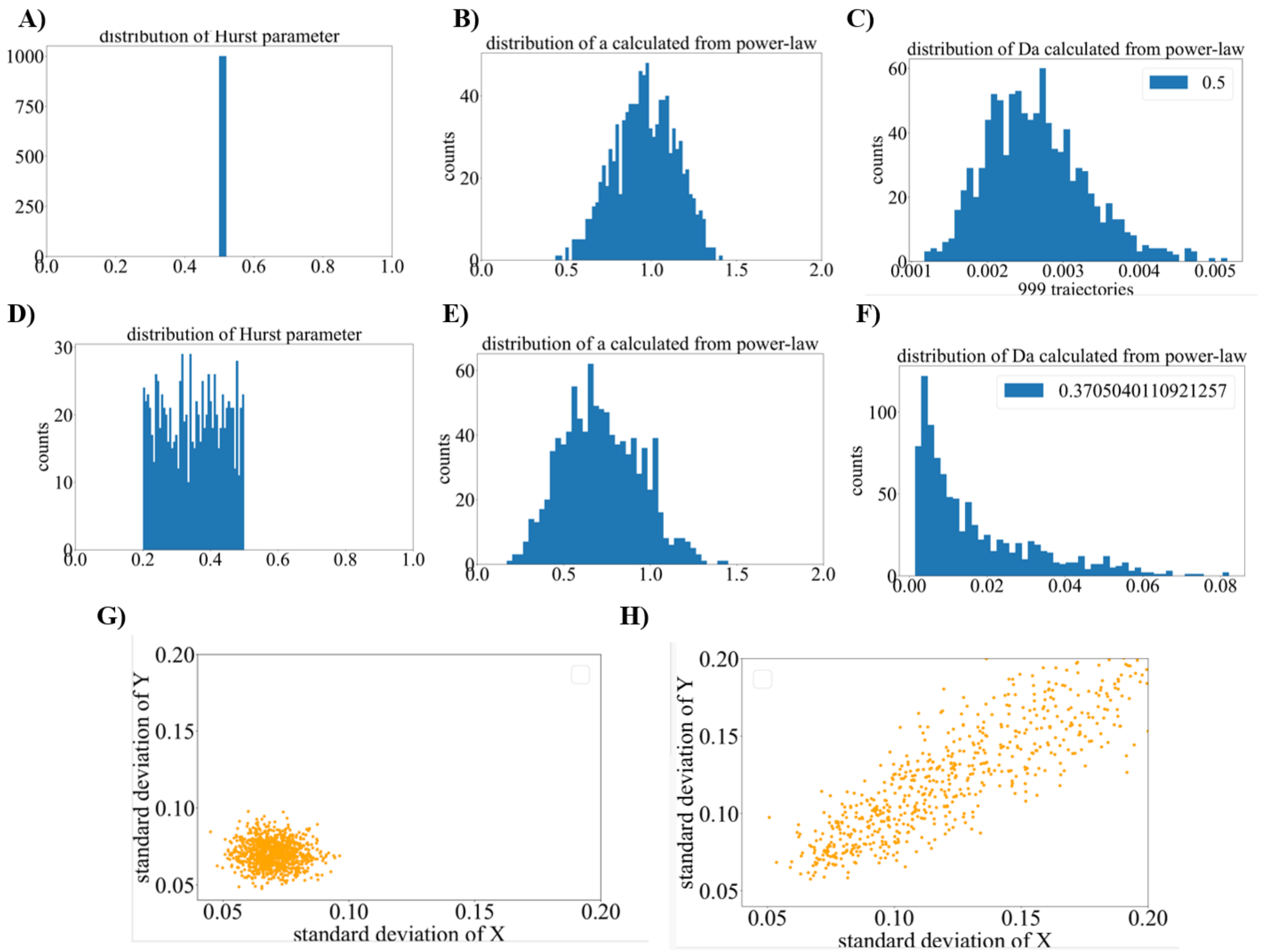


S2. Fractional Brownian motion model implemented in Python. A) A single type of motion in the system (H = 0.5) with corresponding distributions of B) α and C) $D_\alpha$ estimated through the power-law equation (see Methods). D) A mix of motion types (H values ranging from 0.2 to 0.5) with corresponding distributions of E) α\alphaα and F) $D_\alpha$ , also estimated through the power-law equation. Plots of the standard deviation of x and y displacements collected for G) H = 0.5 and H) H values ranging from 0.2 to 0.5.

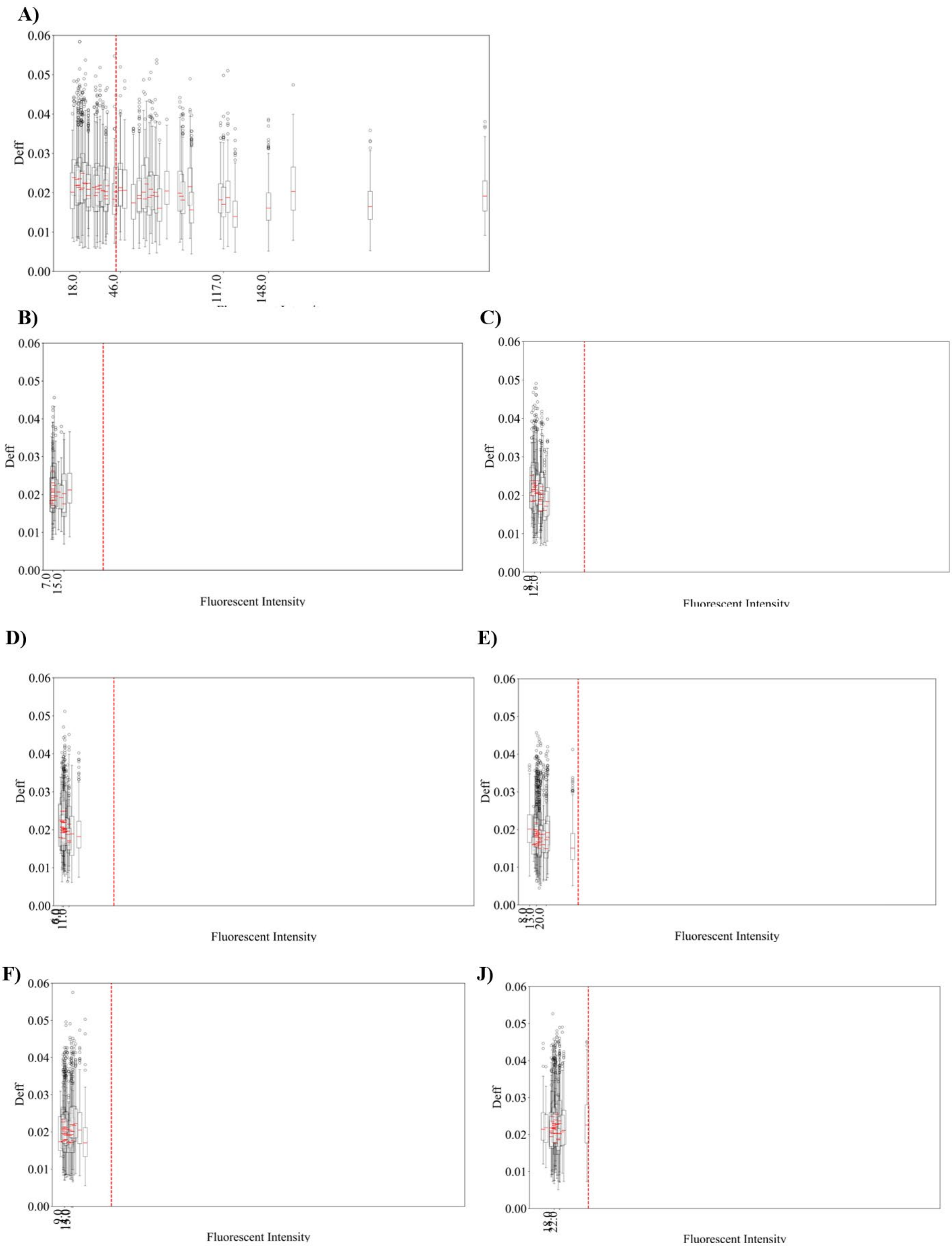
A)
Deff
18.0
46.0
117.0
148.0
B)
Deff
7.0
15.0
Fluorescent Intensity
C)
Deff
12.0
Fluorescent Intensity
D)
Deff
11.0
Fluorescent Intensity
E)
Deff
8.0
13.0
20.0
Fluorescent Intensity
F)
Deff
Fluorescent Intensity
J)
Deff
18.0
22.0
Fluorescent Intensity

Figure S3. Panel of boxplots showing Deff vs. fluorescence intensity within the cell for single tracks analyzed using the power-law fit with $\sigma/\Delta t = 0.5$. A) Colony expressing GEM without expression control; B) Colony expressing GEM under doxycycline treatment at 0.24 µg/mL for 24 hours; C) 0.24 µg/mL for 48 hours; D) 0.5 µg/mL for 24 hours; E) 0.5 µg/mL for 48 hours; F) 1 µg/mL for 24 hours; G) 1 µg/mL for 48 hours.

| Experiment | Statistic | Value | p-value |
|---|---|---|---|
| 0.24 ug/ul dox 2 days | Spearman Correlation | -0.474427663 | 0.005 |
| 0.24 ug/ul dox 2 days | Pearson Correlation | -0.549422006 | 0.001 |
| 0.24 ug/ul dox 1 day | Spearman Correlation | 0.039935171 | 0.871 |
| 0.24 ug/ul dox 1 day | Pearson Correlation | -0.087494548 | 0.722 |
| 0.5 ug/ul dox 2 days | Spearman Correlation | -0.350771892 | 0.039 |
| 0.5 ug/ul dox 2 days | Pearson Correlation | -0.370797777 | 0.028 |
| 0.5 ug/ul dox 1 day | Spearman Correlation | -0.377214265 | 0.025 |
| 0.5 ug/ul dox 1 day | Pearson Correlation | -0.395688889 | 0.019 |
| 1 ug/ul dox 2 days | Spearman Correlation | -0.02133757 | 0.903 |
| 1 ug/ul dox 2 days | Pearson Correlation | 0.040572229 | 0.817 |
| 1 ug/ul dox 1 day | Spearman Correlation | -0.035284369 | 0.841 |
| 1 ug/ul dox 1 day | Pearson Correlation | -0.04760123 | 0.786 |
| GEM | Spearman Correlation | -0.705689463 | 3.04E-10 |
| GEM | Pearson Correlation | -0.558882377 | 3.47E-06 |

**Supplementary Table 1.** Spearman (monotonic change) and Pearson (linear change) correlation coefficients for the dependency of median Deff vs. fluorescence intensity.

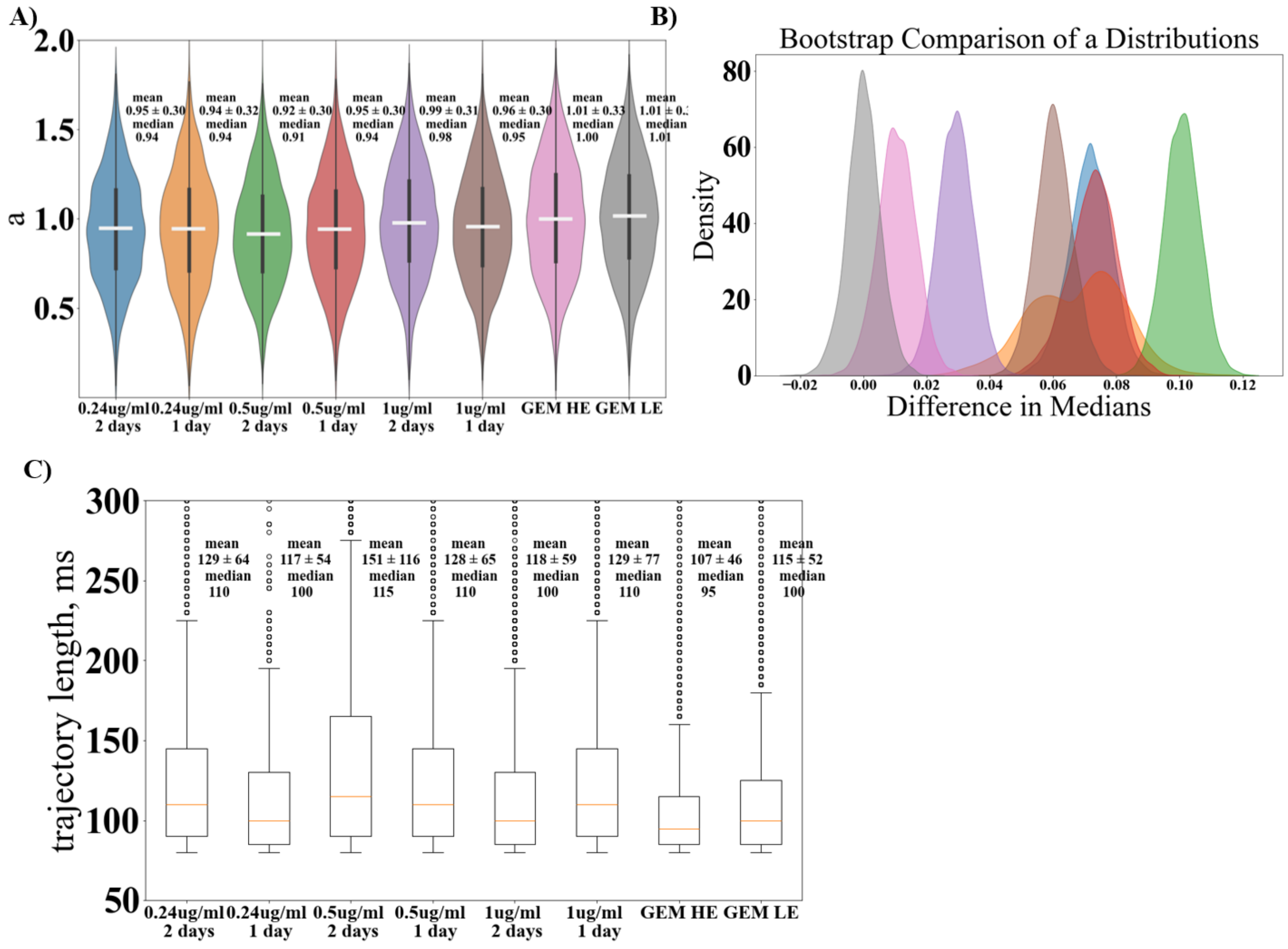


S4. A) (left) Violinplots of $\alpha$ distributions from experiments with different levels of GEM fluorescence (30 cells per group, except 0.24 ug/ml dox 1 day (20 cells) because of low expression) collected from power-law fit with $\sigma/\Delta t = 0.5$. GEM HE = 'High GEM Expression'. GEM LE = 'low GEM Expression'. '*' significant difference with GEM HE estimated by non-parametric bootstrap analysis; (right) The corresponding bootstrap distributions reflecting median differences between groups; C) Boxplots of length in ms for detected trajectories.

Live imaging of cells expressing GEM under a dox-inducible promoter after 0.24 μg/mL doxycycline (Video S1), 0.5 μg/mL doxycycline (Video S2), and 1 μg/mL doxycycline (Video S3), following 24 hours of doxycycline induction. Video S4 shows U2OS cell expressing GEM under a constitutive promoter from the "GEM High Expression" group. Video S5 shows U2OS cell expressing GEM under a constitutive promoter from the "GEM Low Expression" group.
